# An anti-Gn glycoprotein antibody from a convalescent patient potently inhibits the infection of severe fever with thrombocytopenia syndrome virus

**DOI:** 10.1101/434530

**Authors:** Ki Hyun Kim, Jinhee Kim, Meehyun Ko, June Young Chun, Hyori Kim, Seungtaek Kim, Ji-Young Min, Wan Beom Park, Myoung-don Oh, Junho Chung

**Affiliations:** Department of Biochemistry and Molecular Biology, Seoul National University College of Medicine, Seoul, Republic of Korea; Cancer Research Institute, Seoul National University College of Medicine, Seoul, Republic of Korea; Respiratory Virus Laboratory, Institut Pasteur Korea, Gyeonggi-do, Republic of Korea; Department of Internal Medicine, Seoul National University College of Medicine, Seoul, Republic of Korea; Zoonotic Virus Laboratory, Institut Pasteur Korea, Gyeonggi-do, Republic of Korea; Department of Biomedical Science, Seoul National University College of Medicine, Seoul, Republic of Korea

**Author notes:** Asan Institute for Life Sciences, Asan Medical Center, Seoul, Republic of Korea (Hyori Kim). Current affiliation: GlaxoSmithKline, Rockville, Maryland, United States of America (Ji-Young Min). Corresponding author (JC).

## Abstract

Severe fever with thrombocytopenia syndrome (SFTS) is an emerging infectious disease localized to China, Japan, and Korea that is characterized by severe hemorrhage and a high fatality rate. Currently, no specific vaccine or treatment has been approved for this disease. To develop a therapeutic agent for SFTS, we isolated antibodies from a phage-displayed antibody library that was constructed from a patient who recovered from SFTS virus (SFTSV) infection. One antibody, designated as Ab10, was reactive to the Gn envelope glycoprotein of SFTSV and protected host cells and A129 mice from infection in both *in vitro* and *in vivo* experiments. Notably, Ab10 protected 80% of mice, even when injected 5 days after inoculation with a lethal dose of SFTSV. Using cross-linker assisted mass spectrometry and alanine scanning, we located the non-linear epitope of Ab10 on the Gn glycoprotein domain II and an unstructured stem region, suggesting that Ab10 may inhibit a conformational alteration that is critical for cell membrane fusion between the virus and host cell. Ab10 reacted to recombinant Gn glycoprotein in Gangwon/Korea/2012, HB28, and SD4 strains. Additionally, based on its epitope, we predict that Ab10 binds the Gn glycoprotein in 247 of 272 reported SFTSV isolates previously reported. Together, these data suggest that Ab10 has potential to be developed into a therapeutic agent that could protect against more than 90% of reported SFTSV isolates.

**Author summary:** Severe fever with thrombocytopenia syndrome (SFTS) is an emerging infectious disease localized to China, Japan, and Korea. This tick-borne virus has infected more than 5,000 humans with a 6.4% to 20.9% fatality rate. Currently, there are no prophylactic or therapeutic measures against this virus. Historically, antibodies from patients who recovered from viral infection have been used to treat new patients. Until now, one recombinant monoclonal antibody was approved for the prophylaxis of respiratory syntial virus infection. We selected 10 antibodies from a patient who recovered from SFTS and found that one antibody potently inhibited SFTS viral infection in both test tube and animal studies. We determined the binding site of this antibody to SFTS virus, which allowed us to predict that this antibody could bind 247 out of 272 SFTS virus isolates reported up to now. We anticipate that this antibody could be developed into a therapeutic measure against SFTS.

## Introduction

Since its isolation as a novel virus in 2011, cases of the acute infectious disease called severe fever with thrombocytopenia syndrome (SFTS)[1] have risen rapidly in China, Japan, and Korea, posing a risk to public health and increasing the fear of ticks that transmit the deadly SFTS virus (SFTSV). From 2011 to 2016, this emerging tick-borne virus infected 5,360 people in China with an average case fatality rate of 6.40%[2]. After initial reports in 2013 of sporadic SFTS cases in South Korea[3] and Japan[4], South Korea reported 605 cases with an average case fatality of 20.9%[5] and Japan reported 310 cases with an average fatality of 19.4%[6] from 2013 to 2017.

Ticks such as *Haemaphysalis longicornis* and *Rhipicephalus microplus* are implicated as the prominent vectors for transmitting SFTSV[7]. With regards to SFTSV hosts, various vertebrate species are considered to have been infected, as evidenced by high SFTSV seroprevalence in domestic animals in SFTS endemic regions[8,9]. Additionally, reported cases of human-to-human transmission through contact with blood or body fluid, including infections in healthcare workers from patients, pose a further threat to the public[10,11]. Furthermore, the discovery of *H. longicornis* tick in the United States indicates the possibility that SFTSV could spread to other continents, highlighting the need to prevent disease transmission[12].

SFTSV is a single-stranded negative-sense tripartite RNA virus that is classified as a member of the *Phlebovirus* genus, *Phenuiviridae* family, and *Bunyaviriales* order. The genome of SFTSV is comprised of L, M, and S segments, which encode RNA-dependent RNA polymerase (L segment), envelope Gn glycoprotein (M segment), envelope Gc glycoprotein (M segment), nucleoprotein, (S gement) and nonstructural proteins (S segment) [13]. A phylogenetic analysis based on genome sequences of SFTSV isolates found substantial genetic diversity and accumulated mutations, suggesting that SFTSV has existed for decades at minimum[14,15]. However, the difference in virulence between these SFTSV sub-lineages has yet to be determined.

The major clinical features of SFTS include high fever, fatigue, malaise, anorexia, nausea, vomiting, diarrhea, thrombocytopenia, leukocytopenia, and abdominal pain[16,17]. In severe cases, SFTS can include central nervous system manifestations, hemorrhagic signs, and multiple organ dysfunction, which can lead to death[18–21]. No vaccines or therapeutics specific for SFTS have been approved for human use. Recently, a Phase 3 clinical trial of favipiravir (Avigan^®^), an drug approved for the treatment of influenza virus infection in Japan, was initiated to expand its indication to SFTS treatment[22]. Monoclonal antibodies or convalescent sera from SFTS patients were tested to identify potential therapeutic intervention targets, resulting in the identification of SFTSV glycoproteins as molecules required for host cell entry[23,24] and also as critical targets for virus neutralization through the development of humoral immunity. However, generating an antibody for these targets in infected humans was found to be rare, due to the presence of immunodominant decoy epitopes in the nucleoprotein[25], which is a common phenomenon in a pathogenic virus-infected host[26]. But in animal models, the protective effect of human convalescent sera was shown, suggesting that antibody therapy is possible[27]. Thus far, MAb4-5 is the only human neutralizing monoclonal antibody reported, and it was developed using a combinatorial human antibody library from five patients[28]. MAb4-5 binds to domain III of SFTSV Gn glycoprotein[29]. The neutralizing effect of MAb4-5 has been shown only in *in vitro*, and its *in vivo* efficacy remains to be shown.

In this study, we constructed an antibody library from a patient who recovered from SFTS, and selected antibodies against the Gn and Gc glycoproteins. Among these antibodies, Ab10 bound to Gn glycoprotein and showed a potent neutralizing effect both *in vitro* and *in vivo*. In addition, we characterized the conformational epitope of Ab10 using crosslinking coupled mass spectrometry and by testing its reactivity to alanine mutants, which allowed us to estimate the strain coverage of Ab10.

## Results

### Anti-Gn/Gc glycoprotein antibodies were selected from an antibody library generated from a convalescent SFTS patient

In human embryonic kidney (HEK) 293F cell, we produced Gn and Gc glycoproteins fused with a crystallizable fragment of the human immunoglobulin (Ig) heavy chain constant region (Gn-Fc and Gc-Fc) or those fused with the human Ig kappa light chain constant region (Gn-Cκ and Gc-Cκ) and then purified the proteins. We constructed the phage-displayed single-chain variable fragment (scFv) antibody library with a complexity of 1.3 × 10^9^ using peripheral blood mononuclear cells isolated from a patient who had recovered from SFTS. The library was subjected to four rounds of biopanning against either the recombinant Gn-Fc or the Gc-Fc fusion proteins conjugated to paramagnetic beads. We randomly selected phagemid clones from the output titer plate from the last round of biopanning and subjected these clones to phage enzyme-linked immunosorbent assay (ELISA). To minimize the number of clones reactive to the Fc portion of fusion proteins, Gn-Cκ and Gc-Cκ were used as antigens. Positive clones were selected and subjected to Sanger sequencing to determine the scFv nucleotide sequence. We identified five clones reactive to Gn and five clones reactive to Gc. All of these scFv clones were expressed as a scFv antibody fused with Fc (scFv-Fc) in HEK293F cell and purified using affinity chromatography.

### Ab10 mAb potently inhibited the amplification of SFTSV *in vitro*

We tested 10 antibodies for their ability to reduce cytopathic effects (CPE) caused by SFTSV (S1 Fig). One anti-Gn antibody, designated as Ab10, was extremely effective at neutralizing SFTSV by reducing the percentage of cells showing CPE from 90% to 10%. The V_H_ sequences of Ab10 had a 95.9% shared identity with the IGHV3-30*18 germline, excluding the heavy chain complementary determining region (HCDR) 3, whereas the V_κ_ sequence had 86.3% shared identity with the IGKV1-39*01 germline (S2 Fig).

In an immunofluorescence assay (IFA) using Vero cells and an anti-Gn antibody, we determined the proportion of Gn glycoprotein producing cells, which were infected with SFTSV, to measure the neutralizing potency of Ab10. Only 5.6 ± 2.8% (mean ± s.d.) of Vero cells produced Gn glycoprotein when Ab10 was treated at a concentration of 50 μg/ml (956 nM) (Fig 1). Ab10 also showed a dose-dependent protective effect (S3 Fig). When MAb4-5 antibody was applied at the same concentration, 77.8 ± 18.0% of cells produced Gn glycoprotein. When cells were not protected by any antibody, all cells produced Gn glycoprotein and cells not incubated with SFTSV did not produce Gn glycoprotein.

**Fig 1.**
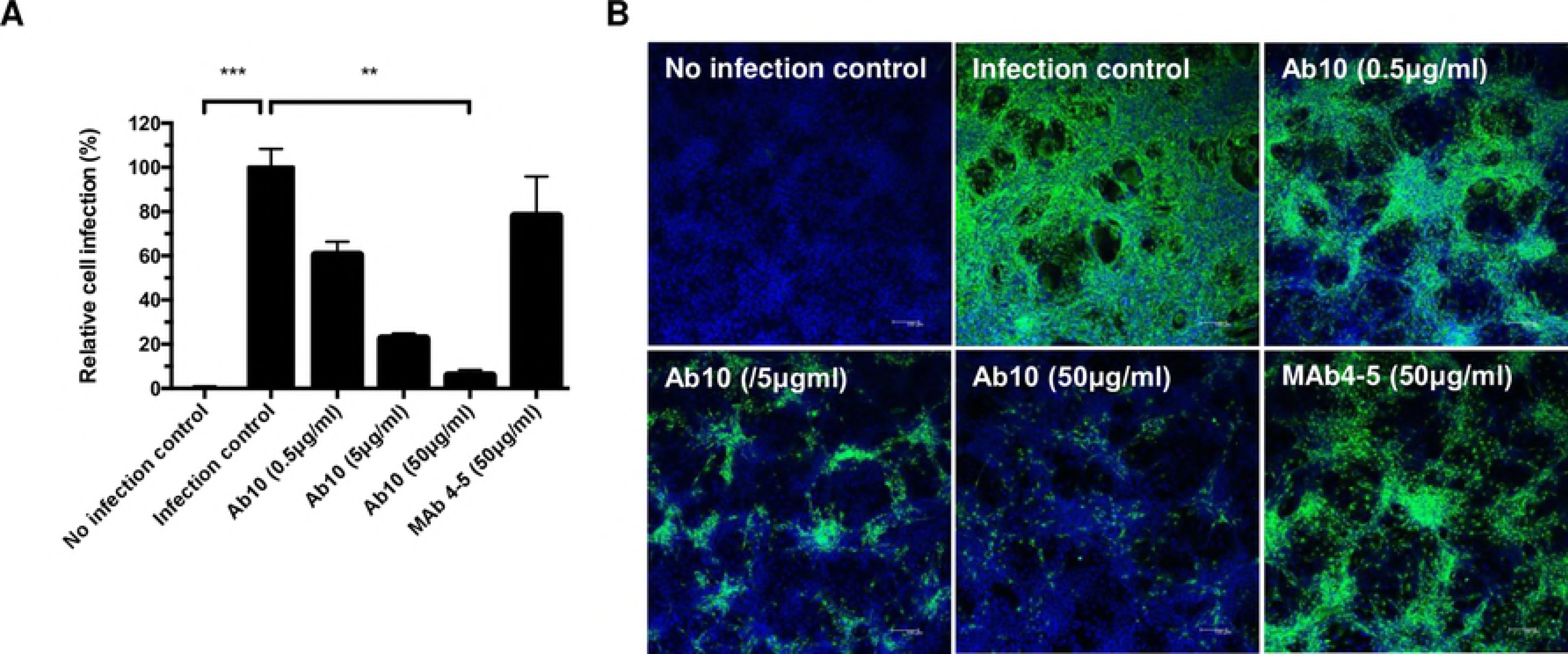
Ab10 has *in vitro* neutralizing activity against Severe Fever with Thrombocytopenia Syndrome virus (SFTSV) To measure neutralizing efficacy, Ab10 scFv-Fc fusion protein was mixed with 100 TCID_50_ of SFTSV (strain: Gangwon/Korea/2012) and added to Vero cells. After incubation for 1 h, the cells were washed and cultured for 2 days. Then, the Gn glycoprotein produced in infected Vero cells was detected in an immunofluorescence assay using anti-SFTSV Gn glycoprotein antibody, with at least five technical replicates. The fluorescence signal intensity of stained SFTSV Gn glycoprotein was used as a quantitative indicator for viral infection. **(A)** The proportion of infected cells compared to non-treated cells was defined as relative cell infection (%) and was plotted. Mab4-5 scFv-Fc fusion protein was also treated in a parallel experiment. Error bars represent standard deviations (s.d.), asterisks indicate a statistically significant difference as determined by a nonparametric Friedman test with a post hoc Dunn’s multiple comparison test (* P ≤ 0.05, ** P ≤ 0.01, *** P ≤ 0.001, **** P ≤ 0.0001). **(B)** Representative images of each treatment group are shown (scale bar, 100 μm). SFTSV Gn glycoprotein and nuclei were stained in FITC (green) and DAPI (blue), respectively.

### Ab10 protected mice from SFTSV infection, even with treatment delayed up to 3 days

Type I interferon (interferon α/β) receptor gene (IFNAR1) deficient A129 mice (n = 5 per group) were subcutaneously injected with the Gangwon/Korea/2012 strain of SFTSV at a dose of either 2 or 20 plaque forming units (PFU). After 1 h, mice were intraperitoneally administered with either phosphate-buffered saline (PBS), Ab10, MAb4-5, or a human IgG_1_ isotope control antibody at a dose of 600 μg (approximately corresponding to 30 mg/kg of body weight); for 4 days at 24 h intervals, the injection of the same amount of antibody was performed (Fig 2A).

**Fig 2.**
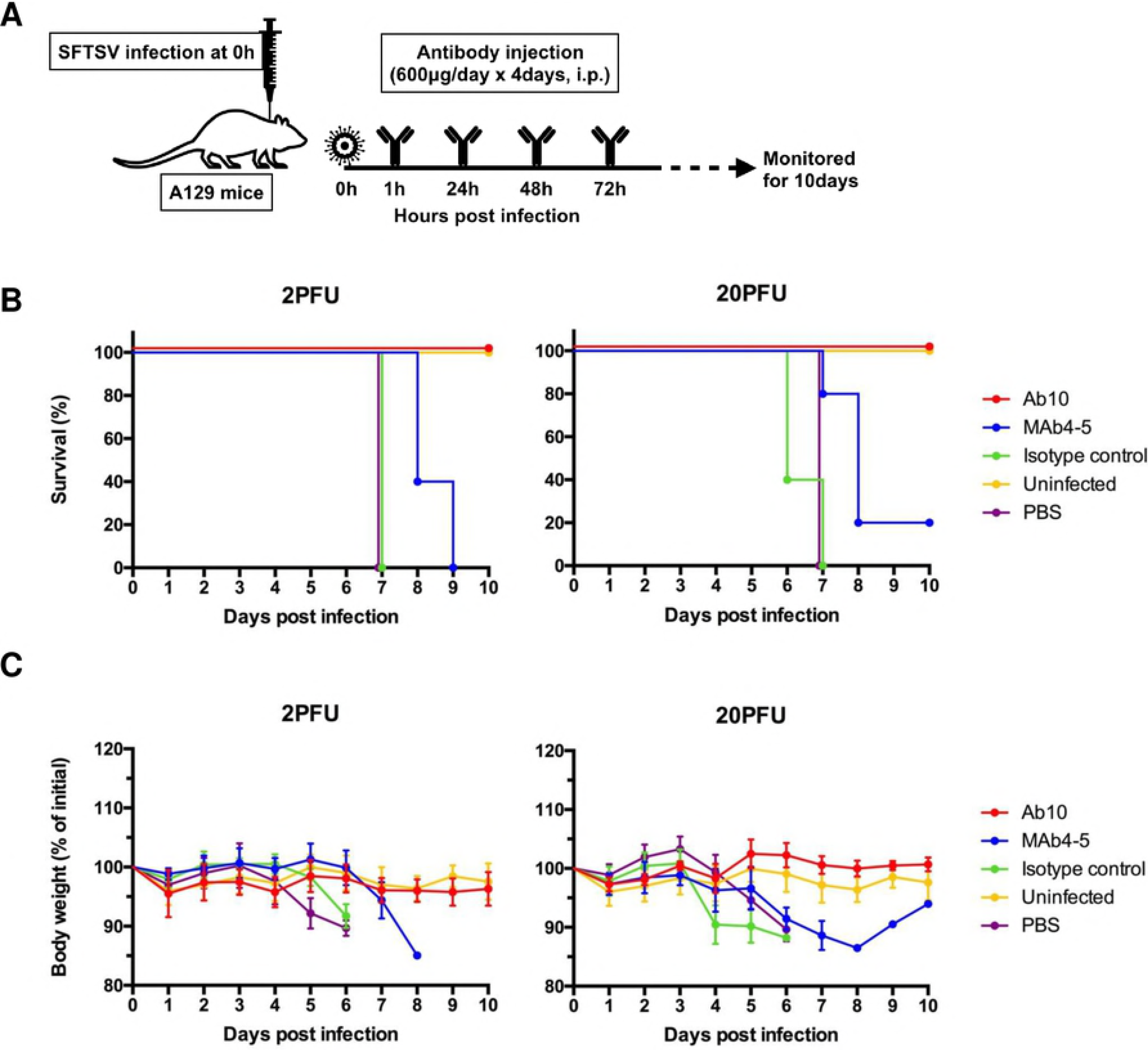
Ab10 protected mice from SFTSV infection. The overall scheme for the administration of virus and antibody is described in **(A)**. Eight-week-old A129 mice (n = 5 per group) were inoculated with 2 or 20 PFU of SFTSV through a subcutaneous route. At 1, 24, 48, and 72 h post infection, infected mice were intraperitoneally administered with 600 μg of Ab10, MAb4-5, IgG_1_ isotype control antibody, or PBS vehicle control. Percentages of survival **(B)** and body weight relative to the day of virus inoculation **(C)** were monitored daily until 10 days post infection. Survival was determined by the Kaplan-Meier method. Relative body weight values in **(C)** are presented as the means with standard deviations of surviving mice in each group.

In the groups treated with PBS or an isotype control antibody, all mice died within 7 days at both viral doses (Fig 2B and 2C). At 4 days post infection (d.p.i.) with a dose of 2 PFU, approximately 10% of body weight was lost; at 3 d.p.i. with 20 PFU, 10–15% of body weight was lost. All mice treated with Ab10 survived both viral doses and did not have any weight loss. With MAb4-5 treatment, death occurred in all mice treated with a 2 PFU viral dose and 80% of mice treated with a 20 PFU dose, and significant weight loss was observed in all these mice.

In the delayed treatment model, the antibody treatment started from 1, 3, 4, or 5 d.p.i. and continued for 4 consecutive days (Fig 3A). At a 2 PFU viral dose, all mice survived when treatments with Ab10 were delayed until 3 d.p.i., and 80% survived when the treatment was delayed until 4 d.p.i. or 5 d.p.i.(Fig 3B and 3C). Mice not treated until 4 d.p.i. had significant weight loss. At a 20 PFU viral dose, delaying Ab10 antibody treatment until 1 or 3 d.p.i. protected all or 80% of mice, respectively. Mice with treatment delayed until 1 d.p.i did not lose weight, whereas mice with treatment delayed until 3 d.p.i. lost 8% of body weight. When treatment was delayed until 4 d.p.i or later, all the mice died.

**Fig 3.**
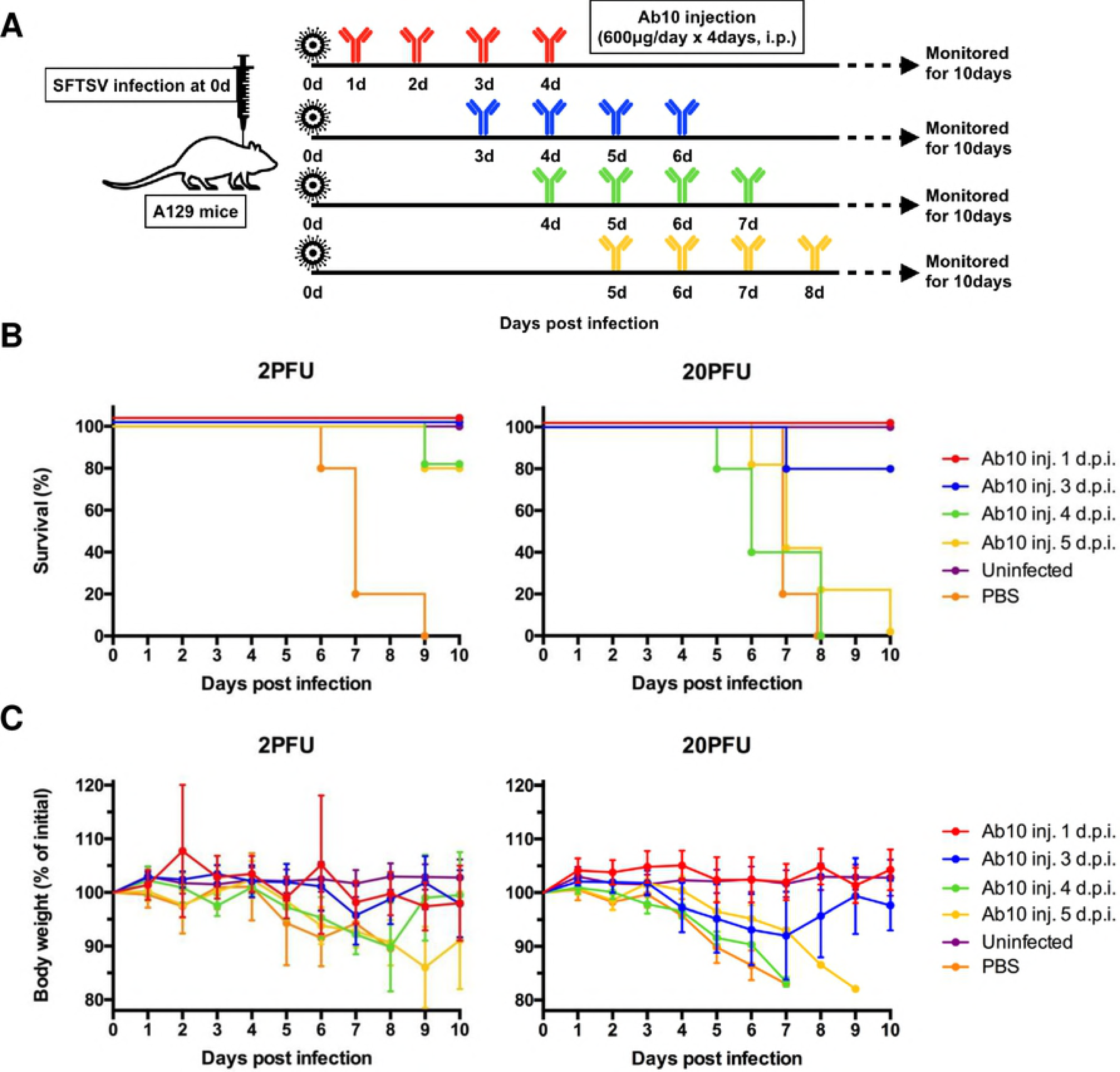
Delayed administration of Ab10 also protected mice from SFTSV infection up to 3 days after inoculation of the virus. The overall scheme for the virus challenge and delayed antibody administration are described in **(A)**. Eight-week-old A129 mice (n = 5 per group) were inoculated with 2 or 20 PFU of SFTSV through a subcutaneous route. From 1, 3, 4, or 5 days post infection, infected mice were intraperitoneally administered with 600 μg of Ab10 per day for 4 consecutive days. Percentages of survival **(B)** and weight relative to the day of virus inoculation **(C)** were monitored daily until 10 days post infection. Survival was determined by the Kaplan-Meier method. The values in **(C)** are presented as the means with standard deviations of surviving mice in each group.

### Ab10 binds to recombinant Gn glycoprotein with high affinity in a broad variety of strains

To check the reactivity of Ab10 to SFTSV strains other than Gangwon/Korea/2012, we overexpressed and purified recombinant Gn glycoproteins of other SFTSV strains. Among the 272 SFTSV strain sequences deposited in the Virus Pathogen Database and Analysis Resource (ViPR), we selected the strains HB29, AH15, SD4, YG1, which belong to other clusters (S4 Fig), to compare their reactivity with well-known virus isolates from China and Japan. We successfully overexpressed Gn glycoprotein from HB29 and SD4 as a Fc fusion protein and subjected these proteins to ELISA. Ab10 IgG_1_ successfully bound to Gn glycoproteins from the HB29 and SD4 strains in a dose-dependent manner, at concentrations ranging from 10 pM to 1 nM (Fig 4A). Further, the amount of antibody bound to the HB29 and SD4 Gn glycoproteins coated on the ELISA plate was higher than that of the Gangwon/Korea/2012 glycoproteins, at most of the tested concentrations. We also found that Mab4-5 was reactive to Gn glycoprotein from the HB29 and SD4 strains (Fig 4B).

**Fig 4.**
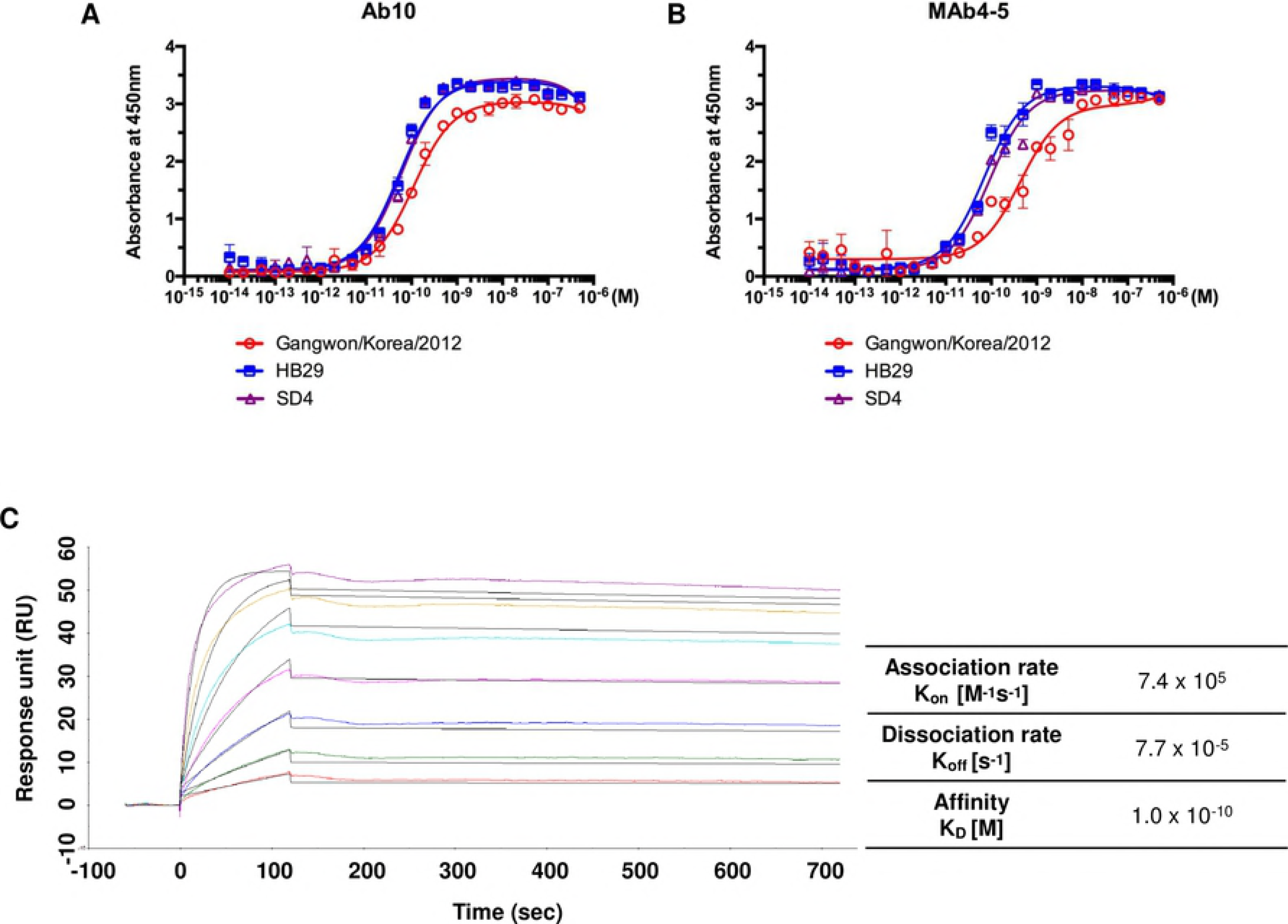
Ab10 also bound to Gn glycoprotein of HB29 and SD4 strains with comparable affinity to that of Gangwon/Korea 2012. Binding properties of human IgG_1_ monoclonal antibody Ab10 **(A)** and MAb4-5 **(B)** to recombinant Gn glycoprotein ectodomain of Gangwon/Korea 2012, HB29, and SD4 strains were measured by enzyme-linked immunosorbent assay (ELISA). Non-linear regression curves were fitted to a one site specific saturation binding model and the mean absorbance at 450 nm with standard deviation (s.d.) error bars are shown at each antibody concentration. **(C)** Surface plasmon analysis of Ab10 antibody was performed on the CM5 chip with an immobilized anti-histidine antibody binding to poly-histidine tagged SFTSV Gn ectodomain. The experimental data at concentrations of 80, 40, 20, 10, 5, 2.5, and 1.25 nM Ab10 antibody are shown in color, and the fitted curves are shown in black. Calculated rate constants are shown in the table.

We used surface plasmon resonance analysis to determine the kinetics of Ab10 binding to the Gn glycoprotein of Gangwon/Korea/2012. Ab10 bound to Gn glycoprotein with an equilibrium dissociation constant (K_D_) of 104 pM and found an association rate (K_*on*_) of 7.4 × 10^5^ M^−1^s^−1^ and a dissociation rate (K_*off*_) of 7.7 × 10^−5^ s^−1^ (Fig 4C).

### Ab10 binds to a non-linear epitope on domain II and the stem region of the Gn glycoprotein

In an immunoblot analysis using recombinant Gn glycoprotein from the Gangwon/Korea/2012 strain, Ab10 did not react to Gn glycoprotein, whereas some other anti-Gn antibodies were reactive (S5 Fig). Based on this observation, we concluded that the antibody reacts to a non-linear epitope.

To discover the site where Ab10 binds, we performed crosslinking coupled mass spectrometry using a deuterium isotope-labeled homo-bifunctional linker, which forms covalent bonds between amino acid residues within the interface of the antibody-antigen complex as described previously[30]. We found that cross-linkers bound to five amino acid residues (318Y, 324R, 326K, 328Y, and 331S) within domain II of the SFTSV Gn glycoprotein and also to four amino acid residues (371K, 372S, 379H, and 383S) within the stem region (Fig 5A).

**Fig 5.**
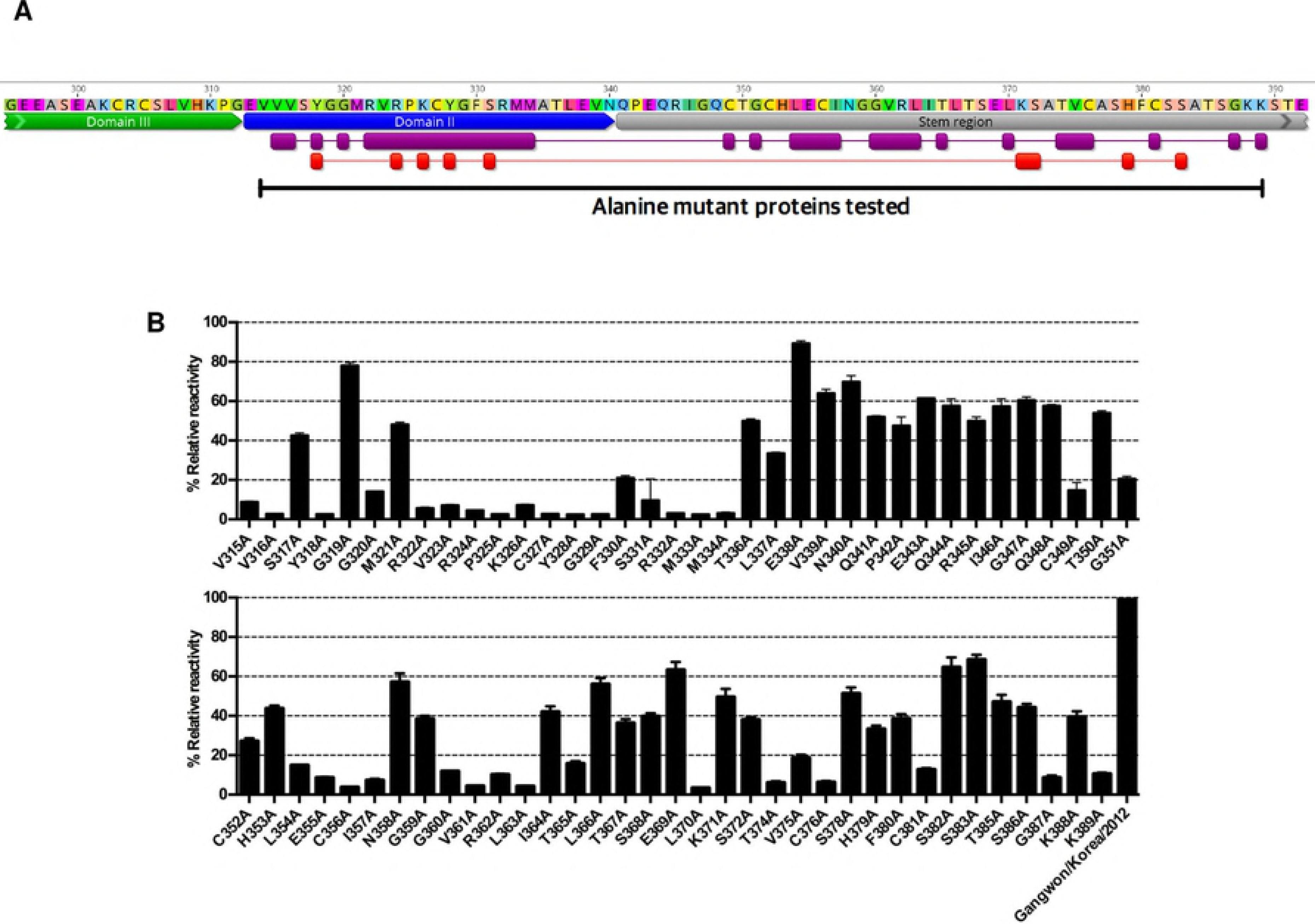
The epitope of Ab10 was determined by alanine mutant analysis. The conformational epitope of Ab10 antibody on Gn glycoprotein ectodomain was determined by measuring antibody binding activity to recombinant mutant proteins with amino acid residues that were substituted with alanine at residues corresponding to 315-389. **(A)** Epitopes predicted by cross-linker assisted mass spectrometry are shown in red, and alanine substituted residues that affected Ab10 antibody binding are shown in purple. The overlapping domain II (blue annotation) and region upstream of the stem region (gray annotation) are also indicated. **(B)** The reactivity of Ab10 to each alanine mutant is represented as relative reactivity, which was calculated using absorbance values (Abs) as follows: % Relative reactivity = [100 x {(Abs of mutant captured by Ab10) / (Abs of mutant captured by HA antibody)} / {(Abs of wildtype captured by Ab10) / (Abs of wildtype captured by HA antibody)}]. Bars indicate the mean and standard deviation (s.d.).

Based on this observation, we prepared several alanine-replacement mutants that spanned from 315V to 389K and tested their reactivity to Ab10 using ELISA. All the mutants were expressed with an HA peptide at the carboxy terminal and the tags were used to measure the relative amount of each mutant. Alanine mutant proteins were captured by the Ab10 antibody, which was coated on the ELISA plate. Then, we measured the amount of captured mutant proteins by detecting the Fc portion of protein. The signals detected by capturing the HA peptide were used to normalize expression of mutant proteins. We measured the reactivity of Ab10 to alanine mutant Gn proteins, relative to wild type Gn glycoprotein, and found that alanine replacement of the amino acid residues in domain II, from V315 to M334, reduced the reactivity of Ab10 by more than 60%, except for S317, G319, and M321. This finding was consistent with our results from the crosslinking coupled mass spectrometry (Fig 5A and 5B).

In the stem region, replacing the cystine residues (C349, C356, C376, and C381) reduced the relative reactivity by more than 80%. This observation was consistent with a previous report that the structural stability of Gn was disrupted by a C356A mutation[29]. Also, mutation of the flanking residues of cystine, corresponding to G351, L354, E355, I357, T374, and V375 also reduced the reactivity by over 60%. In the cases of mutation residues which are distant from cysteine residues, mutation of G360, V361, R362, L363, T365, L370, G387, and K389 residues reduced the relative reactivity by more than 80%. The other residues had minor effects on reactivity. Overall, Ab10 binding to Gn was predicted to be affected by 25 amino acid residues within domain II and the stem region of SFTSV Gn glycoprotein. Because Gn glycoproteins from 247 isolates have conserved sequences for these 25 amino acid residues, we expect that Ab10 can react with 90.8% (247 out of 272 isolates) of SFTSV isolates currently reported (S6 Fig).

## Discussion

Antibodies play a pivotal role in preventing viral entry into cells and can kill infected cells through antibody-dependent cellular cytotoxicity or complement-dependent cytotoxicity[31–33]. Polysera from recovered patients or from vaccinated donors have been used as prophylactic agents for various viral diseases, including hepatitis B and rabies[34]. As an alternative approach, monoclonal antibodies have also been developed and tested as therapies or prophylaxis for viral diseases. Palivizumab (Synagis^®^) was market-approved for the prophylaxis of respiratory syncytial virus (RSV) in 1998. Antibodies against HIV[35–37], RSV[38], Ebola virus[39], and influenza virus[40,41] demonstrated potent efficacy in animal models. Antibodies targeting emerging or re-emerging viruses, including MERS-CoV[42–44] and Zika virus[45–47], were also developed and are being tested in clinical trials. In the past several decades, antibodies have become the one of the major therapeutic agents for cancer and autoimmune disease with indications that have rapidly broadened in recent years. The recent technical improvements in the discovery and manufacturing steps of therapeutic antibody production have also allowed rapid and successful antibody development to combat emerging infectious diseases[48].

Until now, SFTS patients have been reported from China, South Korea, and Japan, and the number of patients has increased each year[2,5,6]. However, SFTS fatality varies among the three countries[21]. The average case fatality rate in China from 2011 to 2016 was 6.40%[2]. Those in South Korea and Japan after 2013 were much higher; 20.9%[5] or 19.4%[6], respectively. In the Virus Pathogen Database and Analysis Resource (ViPR), 272 sequences of SFTSV isolates are currently deposited. But it is unknown if there is any significant variability in the virulence of these isolates. Previous reports showed that mice died 5 to 7 days after infection with 10^6^ focus forming units (FFU) of the YG1 strain[27] or 10^6^ TCID_50_ of the SPL010 strain[49]. Based on these observations, we first inoculated A129 mice with 2 × 10^5^ PFU of the Gangwon/Korea/2012 strain and observed that all mice died 4 days after infection. With a 20 PFU dose of the Gangwon/Korea/2012 strain, mice died 5 to 7 days after infection.

We also observed that the amount of body weight losses at death were much higher in our study than those in other studies.[49]. In our data, A129 mice died after losing 15% of their body weight. But in the SPL010 strain, mice died after losing 30% of their body weight. These results might be due to a difference in virulence between the strains. Differences in virulence between strains of the Rift Valley fever virus (RVFV), a phlebovirus similar to SFTSV, have been reported[50]. Or it might be due to differences in the animals tested, because STAT2 knock out Syrian hamsters challenged with 10 PFU of the HB29 strain showed a similar fatality rate[51].

The mechanisms of antibody inhibition of viral replication inside host cells have been studied extensively, especially in the case of influenza virus. The most-widely known mechanism is for an antibody to bind the portion of virus that interacts with the host cell receptor, thereby blocking the interaction between the virus and the host cell[52]. Another group of antibodies was reported to bind the stem region of influenza hemagglutinin that is critical for conformational rearrangements that occur during membrane fusion[53–55]. This mechanism has more potential to be utilized for clinical development, because the stem region has fewer mutations than the receptor binding site. Additionally, several groups, including ours, have elucidated unconventional virus neutralizing mechanisms that affect the infection steps that occur after membrane fusion[56,57].

In our crosslinking coupled mass spectrometry and alanine mutant studies, the Ab10 epitope was confined to domain II and the stem region of the Gn glycoprotein. Although the crystal structure of the phlebovirus Gn glycoprotein stem region has not yet been solved, a recent report showed a cryo-electron microscopy map of RVFV, and depicted the crystal structure of the RVFV Gn glycoprotein head region without a stem region[58]. The report also describes the membrane fusion mechanism of RVFV that is mediated by a low pH induced exposure of the hydrophobic Gc fusion loop. At a neutral pH, the Gn domain II (β-ribbon domain) shields the Gc fusion loop in the pre-fusion state and prevents premature fusion. Based on this report, we hypothesize that Ab10 simultaneously binds to domain II and the stem region of the Gn glycoprotein and prevents un-shielding of the Gc fusion loop.

In conclusion, Ab10 is a monoclonal antibody that has shown therapeutic efficacy in a mouse SFTSV infection model. Although the neutralization efficacy of Ab10 was only tested in the Gangwon/Korea/2012 strain that was cultured in Vero cells, we confirmed its binding capability to recombinant SFTSV Gn in the HB29 and SD4 strains, which are both from China. According to the epitope revealed in this study, Ab10 is estimated to interact with the majority of SFTSV isolates currently reported. Based on these results, we believe that Ab10 has sufficient potential to be developed as a prophylactic and therapeutic agent for a broad variety of SFTS isolates.

## Materials and methods

### Ethics statements: human subjects and animal models

The studies involving recovered patient’s blood samples were reviewed and approved by the Institutional Ethics Review Board of Seoul National University Hospital (IRB approval number: 1405-031-576). All of the patients were adults and submitted written informed consent. All animal studies were conducted in an Animal Biosafety Level 3 (ABSL-3) facility at the Institut Pasteur Korea according to the principles established by the Animal Protection Act and the Laboratory Animal Act in Republic of Korea. Interferon α/β receptor knockout (IFNAR1^−/−^, A129) mice (B&K Universal, Hull, UK) were bred, raised, and genotyped at Institut Pasteur Korea. All experimental procedures were reviewed and approved by the Institutional Animal Care and Use Committee at the Institut Pasteur Korea (Animal protocol number: IPK-17003-1).

### Production of recombinant SFTSV Gn/Gc glycoprotein fusion proteins

The SFTSV Gn glycoprotein amino acid sequences of various isolates used in this study were retrieved from the Virus Pathogen Database and Analysis Resource (ViPR). To obtain SFTSV Gn glycoprotein ectodomain coding DNA strands, human codon optimized DNA sequences corresponding to amino acid sequences from 20 to 452 of GenBank Accession No. ADZ04471 (Strain HB29), ADZ04477 (Strain SD4), ADZ04486 (Strain AH 15), BAN58185 (Strain YG1), AGT98506 (Strain Gangwon/Korea/2012) were synthesized (GenScript, Piscataway, NJ, USA and Integrated DNA Technologies, Coralville, IA, USA). Human codon optimized DNA sequence of SFTSV Gc ectodomain of strain Gangwon/Korea/2012, corresponding to the sequence from 563 to 1035 of AGT98506, was also synthesized. For the overexpression and purification of recombinant SFTSV Gn / Gc glycoprotein ecotodomain fused to the Fc region of human immunoglobulin heavy constant gamma1 (IGHG1), termed Gn-Fc / Gc-Fc, or fused to the human immunoglobulin kappa constant region (IGKC), termed Gn-Cκ / Gn-Cκ, SFTSV Gn / Gc glycoprotein ectodomain encoding genes were cloned into the modified pCEP4 vector (V04450, Invitrogen, Carlsbad, CA, USA) with a leader sequence of the human immunoglobulin kappa chain, two *Sfi*l restriction enzyme sites, and the Fc region of human IGHG1 or human immunoglobulin kappa constant region, as previously described[59,60]. Subsequently, the vectors were used to transfect HEK 293F (R79007, Invitrogen) or Expi293F cells (A14527, Invitrogen) using polyethylenimine (23966-1, Polysciences, Warrington, PA, USA), then the transfected cells were cultured in FreeStyle™ 293 expression medium (12338026, Gibco, Thermo Fisher Scientific, Waltham, MA, USA). Overexpressed recombinant SFTSV Gn and Gc glycoprotein fusion proteins were purified by affinity chromatography using MabSelect™ or KappaSelect™ columns with the ÄKTA Pure chromatography system (11003495, 17545811, 29018225, GE Healthcare, Chicago, IL, USA), following the protocol provided by the manufacturer.

For alanine-scanning mutagenesis, SFTSV Gn glycoprotein with amino acid residues (315-389) substituted with alanine were produced by cloning synthesized DNA fragments (Integrated DNA Technologies) into modified pCEP4 vector, as described above. Subsequently, influenza hemagglutinin (HA) tag sequence (YPYDVPDYA) was introduced to the C-terminus of the Fc region of human immunoglobulin heavy gamma1 and the whole protein, designated as Gn-Fc-HA, was produced as described above.

In order to produce histidine tagged SFTSV Gn glycoprotein, a ligand for surface plasmon resonance analysis, a Gn-Cκ with six carboxy-terminal poly-histidine residues was designed and produced as described above.

### Human antibody library construction and antibody selection

Peripheral blood mononuclear cells of a patient who recovered from SFTS were collected using a Ficoll-Paque density gradient medium (17144002, GE Healthcare). Total RNA was isolated using TRIzol Reagent (15596018, Invitrogen), and cDNA was synthesized using a SuperScript III first-strand cDNA synthesis kit with oligo dT priming (18080051, Invitrogen). From this cDNA, a phage-display library of human single-chain variable fragments (scFv) was constructed, and four rounds of biopanning were performed to select scFv antibody clones from the library, as previously described[61,62]. For each round of biopanning, recombinant SFTSV Gn-Fc coated onto paramagnetic Dynabeads (14302D, Invitrogen) were used. To select SFTSV glycoprotein binding clones, phage ELISA was performed as previously described, using Gn or Gc glycoprotein-coated microtiter plates, scFv displaying phages, and horseradish peroxidase (HRP) conjugated anti-M13 antibody (11973-MM05, Sino Biological, Beijing, China)[62]. The nucleotide sequences of positive scFv clones were determined by Sanger nucleotide sequencing (Cosmogenetech, South Korea). Germline sequences of selected antibody variable regions were analyzed by the National Center for Biotechnology Information (NCBI) IgBLAST.

### Production of single-chain variable fragment antibodies and IgG_1_ antibodies against SFTSV Gn glycoprotein

The genes encoding the variable heavy chain and variable light chain of Ab10 and MAb4-5[28] were synthesized (Integrated DNA Technologies, GenScript) and fused with human heavy chain constant region gene (IgG_1_) and human kappa light chain gene, and then cloned into an eukaryotic expression vector, as described previously[63,64]. The expression vectors were transfected into HEK 293F cells. The IgG_1_ molecule was purified from the culture supernatant by affinity chromatography using MabSelect™ as described above. Genes encoding the scFv-Fc fusion protein and the scFv-Cκ fusion protein were synthesized and cloned into a pCEP4 vector (Invitrogen). After transfection into HEK 293F cells, the recombinant proteins were overexpressed and purified as described above.

### SFTSV preparation and immunofluorescent imaging-based neutralization test

The SFTSV strain of Gangwon/Korea/2012[3] was propagated in Vero cells (10081, Korean Cell Link Bank) with Roswell Park Memorial Institute (RPMI)-1640 medium (LM 011-01, Welgene, Daegu, South Korea) supplemented with 2% heat-inactivated fetal bovine serum (16000044, Gibco) and penicillin-streptomycin (10378016, Gibco). The fifty-percent tissue culture infective dose (TCID_50_) values were titrated on Vero cells using the Reed-Muench method[65]. Ab10 or MAB4-5 scFv-Fc fusion protein was serially diluted in 10-fold increments from a 50 μg/ml concentration, then mixed with an equal volume of 100 TCID_50_ SFTSV, and incubated at 37°C for 1 h. The virus-antibody mixture was transferred onto Vero cells in 8-well chamber slides (154534, Thermo Scientific, Waltham, MA, USA) and incubated at 37°C for 1 h. For the no infection control group, no virus was added to cells. In contrast, for the infection control group, no antibody was incubated with virus. After removing the virus-antibody mixture, cells were cultured for 2 days. For the IFA, cultured cells were fixed with 4% paraformaldehyde in PBS for 1 h at room temperature. Slides were blocked and permeabilized with PBS containing 0.1% Triton X-100 and 1% bovine serum albumin, followed by incubation with 5 μg/ml of anti-SFTSV Gn glycoprotein antibody[66] at 4°C overnight. After washing, cells were incubated for 1 h at room temperature with 1:100 diluted fluorescein isothiocyanate (FITC)-conjugated anti-rabbit IgG Fc antibody (111-095-046, Jackson ImmunoResearch, West Grove, PA, USA). To stain the nucleus, 4’,6-Diamidino-2-phenylindole dihydrochloride (DAPI) was used. Fluorescence image of cells was monitored under a confocal laser scanning microscope (TCS SP8, Leica, Wetzlar, Germany).

### *In vivo* efficacy test

For animal experiments, the titer of SFTSV was measured by plaque forming assay[67]. Either 2 or 20 plaque forming units (PFU) of Gangwon/Korea/2012 strain SFTSV in 200 μl of PBS were inoculated in 8- to 10-week-old male or female A129 mice by a subcutaneous (s.c.) injection route. After an hour of infection, mice were administered with Ab10 IgG_1_ antibody or a PBS vehicle control through an intraperitoneal (i.p.) injection route, at 30 mg/kg of body weight for every 24 h for a consecutive 4 days. Palivizumab (MedImmune, Gaithersburg, MD, USA) or Mab4-5 IgG_1_ was used as an isotype control or a positive control antibody, respectively. In the delayed treatment model, the infected mice were treated with antibodies at 1, 3, 4, or 5 days post infection (d.p.i.) for 4 days consecutively. Body weight and survival of mice were monitored until 10 days post infection.

### Enzyme-linked immunosorbent assays

In order to measure the binding activities of the Ab10 and MAb4-5 IgG_1_ antibodies, 96-well half-area microplates (3690, Corning, Corning, NY, USA) were coated with Gn-Fc fusion protein and incubated at 4°C overnight. Plates were blocked with 3% skim milk in PBS for 1 h at room temperature. The plates were then washed with PBS and received antibodies that were 10-fold serially diluted from 1 μM to 10 μM in blocking buffer. The plates were then incubated for 2 h at room temperature and washed three times with 0.05% Tween20 in PBS solution. Then, 50 μl of HRP-conjugated anti-human Ig kappa light chain antibody (AP502P, Chemicon, Temecula, CA, USA) diluted in blocking buffer (1:5000) was added into each well. Then, plates were incubated for 1 h at room temperature. After washing, each well received 50 μl of 3,3’,5,5’-Tetramethylbenzidine (TMB) substrate solution (34028, Thermo Scientific). The coloring reaction was stopped by adding 50 μl of 2 M sulfuric acid. The absorbance of each well was measured at 450 nm using a microplate spectrophotometer (Multiskan GO, Thermo Scientific).

### Surface plasmon resonance analysis of Ab10

The kinetics of Ab10 and Gn glycoprotein binding were measured by surface plasmon resonance analysis, using a Biacore T200 instrument with sensor chip CM5, amine coupling kit, and his capture kit (28975001, 29149603, BR100050, 28995056, GE Healthcare). We followed the recommended manufacturer’s protocol for the procedures and conditions of reaction buffers, flow times, flow rates, and concentration of analytes. Briefly, anti-histidine antibody was immobilized on an activated CM5 chip, followed by a deactivation step. Then, histidine tagged Gn-Cº was injected over the flow cells prior to antibody injection. For the association step, all of the Ab10 IgG_1_ antibody in PBS at concentrations of two-fold increments ranging from 1.25 nM to 80 nM was injected for 3 min. For the dissociation step, PBS containing 0.005% of Tween20 was injected for 5 min. After each dissociation step, chip regeneration was performed.

### Conformational epitope mapping by crosslinking coupled mass spectrometry

The epitope of Ab10 antibody was first determined by analyzing the complex of Ab10 antibody and SFTSV Gn-Cκ antigen linked with deuterated cross-linkers, as previously described[30]. Briefly, antibody, antigen, and antibody/antigen complex were characterized by the high mass matrix-assisted laser desorption/ionization (MALDI) mass spectrometry using a MALDI ToF/ToF tandem mass spectrometer (Autoflex III, Bruker, Billerica, MA, USA) equipped with an interaction module (HM4, CovalX, Zürich, Switzerland). Afterwards, the antibody/antigen complex was crosslinked with DSS d0/d12 isotope-labeled homobifunctional N-hydroxysuccinimide esters, followed by reduction alkylation using dithiothreitol, iodoacetamide, and urea. To digest the reduced complex, a proteolytic buffer composed of trypsin, chymotrypsin, endoproteinase Asp-N, elastase, and thermolysin was used. The sample was then analyzed by nano-liquid chromatography (Ultimate 3000, Dionex, Sunnyvale, CA, USA) and Orbitrap mass spectrometry (Q Exactive Hybrid Quadrupole-Orbitrap, Thermo Scientific).

### ELISA for epitope mapping

To measure the binding activities of Ab10 to mutated Gn, Ab10 scFv-Cκ antibody and an anti-influenza virus hemagglutinin antibody (clone 12CA5, Bio X Cell, Lebanon, NH, USA) were coated on a microplate in parallel. Then, plates were blocked with 3% skim milk in PBS for 1 h at room temperature. Transiently transfected supernatant containing recombinant Gn-Fc-HA proteins with alanine substitution was added to each well. After incubation for 2 h at room temperature, the microplate was washed three times with 0.05% Tween20 in PBS solution. Then, HRP-conjugated anti-human IgG Fc antibody (31423, Invitrogen) diluted in blocking buffer was added to each well. The plate was incubated for 1 h at room temperature. After washing, each well received 50 μl of 3,3’,5,5’-Tetramethylbenzidine (TMB) substrate solution (34028, Thermo Scientific). The coloring reaction was stopped by adding 50 μl of 2 M sulfuric acid. The absorbance of each well was measured at 450 nm using a microplate spectrophotometer (Multiskan GO, Thermo Scientific). Relative reactivity was calculated using absorbance values (Abs) as follows: % Relative reactivity = [100 x {(Abs of mutant captured by Ab10) / (Abs of mutant captured by HA antibody)} / {(Abs of wildtype captured by Ab10) / (Abs of wildtype captured by HA antibody)}].

## Data analysis

ELISA and IFA data, including statistical comparisons, were analyzed and graphed using GraphPad Prism software. Fluorescent signal measured by confocal microscope was quantified using Leica Application Suite Advanced Fluorescence software. Mass spectrometry data were analyzed using XQuest and Stavrox software. Plasmon surface resonance data were analyzed using BIAevaluation software. Visualization, alignment, and phylogenic analysis of amino acid sequences were performed with Geneious software.

## Acknowledgements

We thank Myung Jin Lee and Su Jin Choi for setting up the initial experimental conditions for the virus neutralization assay.

## Supporting information

**S1 Fig.**
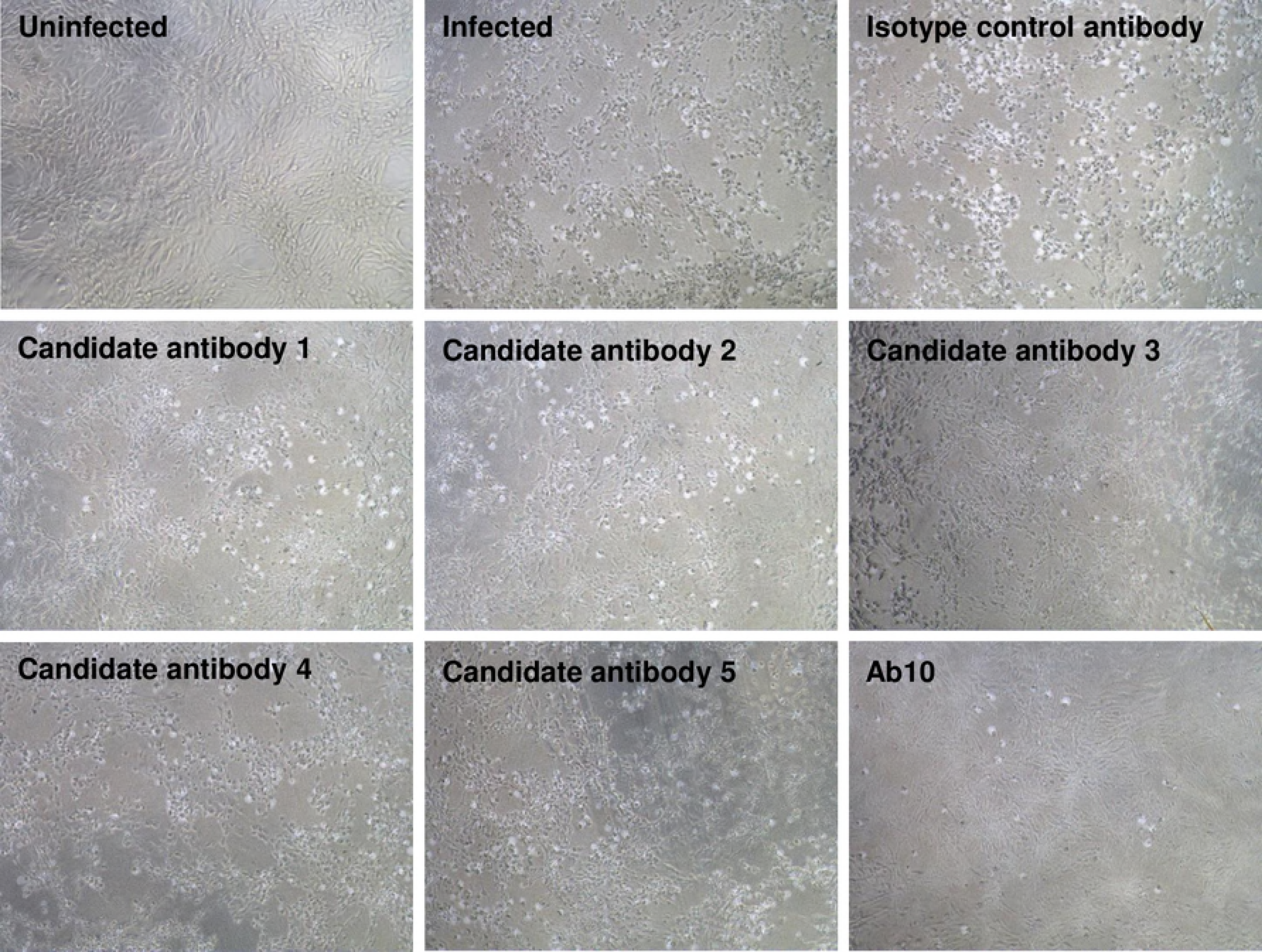
Inhibition of cytopathic effect by SFTSV. The cytopathic effects (CPE) of SFTSV on Vero cells were monitored to evaluate the protective effect of antibody clones. Vero cells at 80% confluency grown in 96-well tissue culture plates were exposed to 100 μl of SFTSV-antibody mixture, which was prepared by mixing 100 TCID_50_/ml SFTSV (Strain: Gangwon/Korea/2012) and 100 μg/ml candidate antibody (scFv-Fc format) at the 1:1 volumetric ratio, and was then pre-incubated for 1 h. After incubating the SFTSV-antibody mixture with cells for 1 h, cells were washed with PBS followed by addition of fresh growth medium for 96 h. Cells were observed under a microscope to evaluate CPE. In control groups, cells not incubated with virus (Uninfected), cells infected without antibody treatment (Infected), cells incubated with virus, and the isotype control antibody (Isotype control antibody) were employed.

**S2 Fig.**
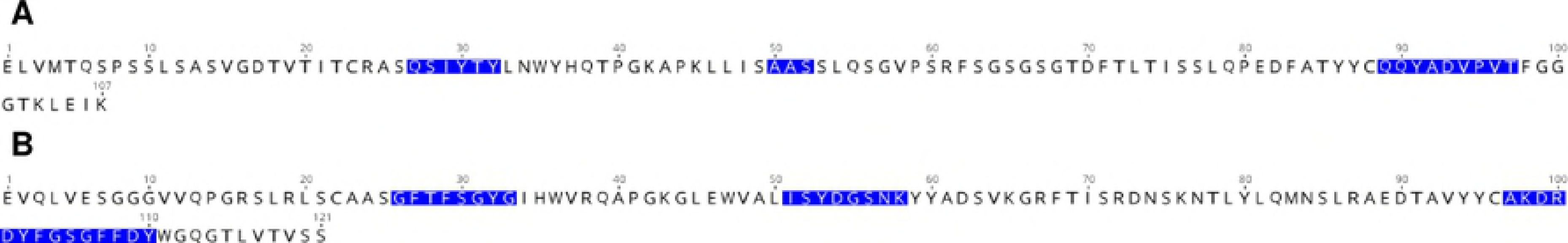
Amino acid sequences of Ab10 antibody variable region. The amino acid sequence of light chain variable region **(A)** and heavy chain variable region **(B)** are shown. Blue letters indicate complementary determining regions (CDR) of each variable region defined by the International Immunogenetics Information System (IMGT).

**S3 Fig.**
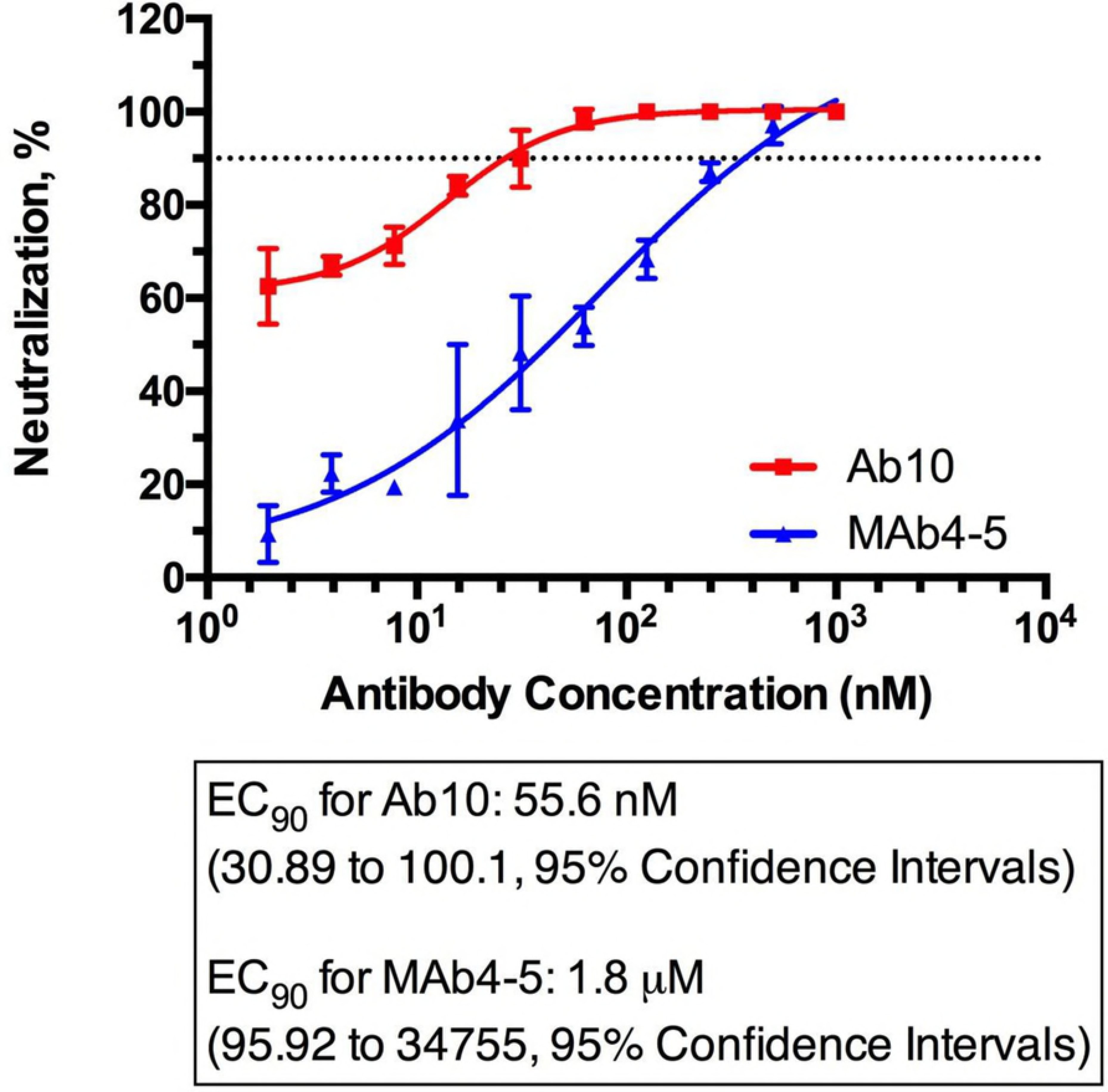
Focus forming assay using Ab10. Thirty to 50 focus forming units (FFU) of SFTSV were incubated with Ab10 scFv-Fc fusion protein at concentrations from 1.95 to 1,000 nM for 1 h at 371C and added to Vero cells in a 24-well tissue culture plate. After incubation for 1 h at 37CC in a 5% CO_2_ incubator, cells were overlaid with 1% methylcellulose in culture medium RPMI with 2% FBS and cultured for 2 days. For detection of SFTSV localized clusters (foci), cells were fixed with a 10% formalin solution for 1 h and incubated with 5 μg/ml of anti-SFTSV Gn glycoprotein detection antibody. After the washing step, cells were incubated for 1 h at room temperature with 1:100 diluted fluorescein isothiocyanate (FITC)-conjugated anti-rabbit IgG Fc antibody (111-095-046, Jackson ImmunoResearch). FFU were determined by counting visible foci under an automated inverted fluorescence microscope (DMI4000 B, Leica). Neutralization efficacy was calculated as the % decreased fraction in the number of foci compared to that of the isotype antibody control group. Dose-response curves were drawn by non-linear regression analysis and fitted to the EC90 (variable slope model). A Mab4-5 scFv-Fc fusion protein was used as a positive control.

**S4 Fig.**
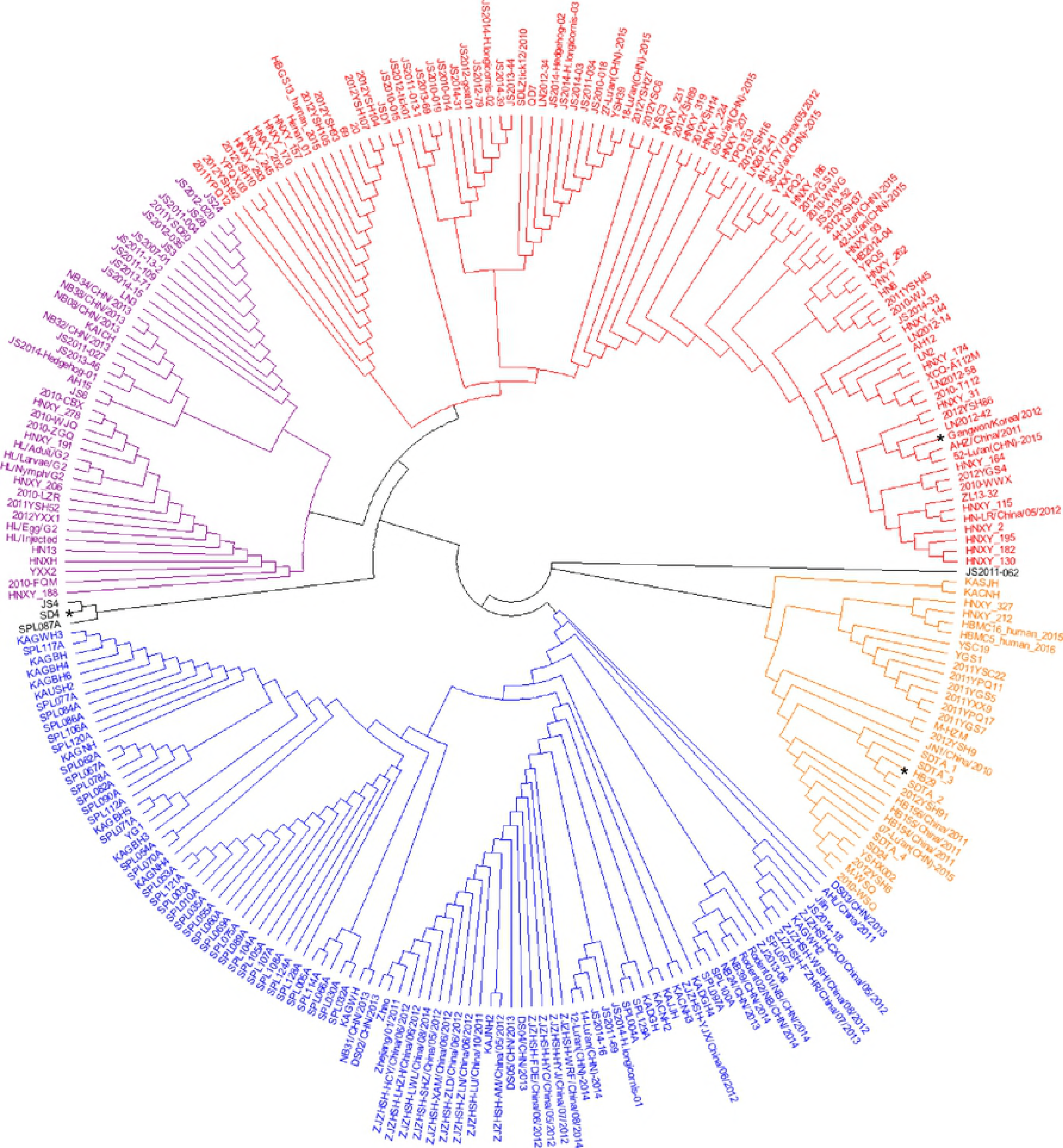
Phylogenetic analysis of SFTSV Gn glycoprotein ectodomain. The amino acid sequence of Gn glycoprotein from 272 SFTSV isolates deposited in ViPR were used for analysis. The sequences were trimmed to retain the amino acid residues from 20-452 that corresponded to the ectodomain. Trimmed sequences were analyzed, and a phylogenetic tree was built in a circular tree layout using the neighbor-joining method with a Jukes-Cantor genetic distance model. The names of isolates are labeled beside the tip of each branch. Asterisks at the tip of branches indicate the isolates that were tested for binding activity of Ab10.

**S5 Fig.**
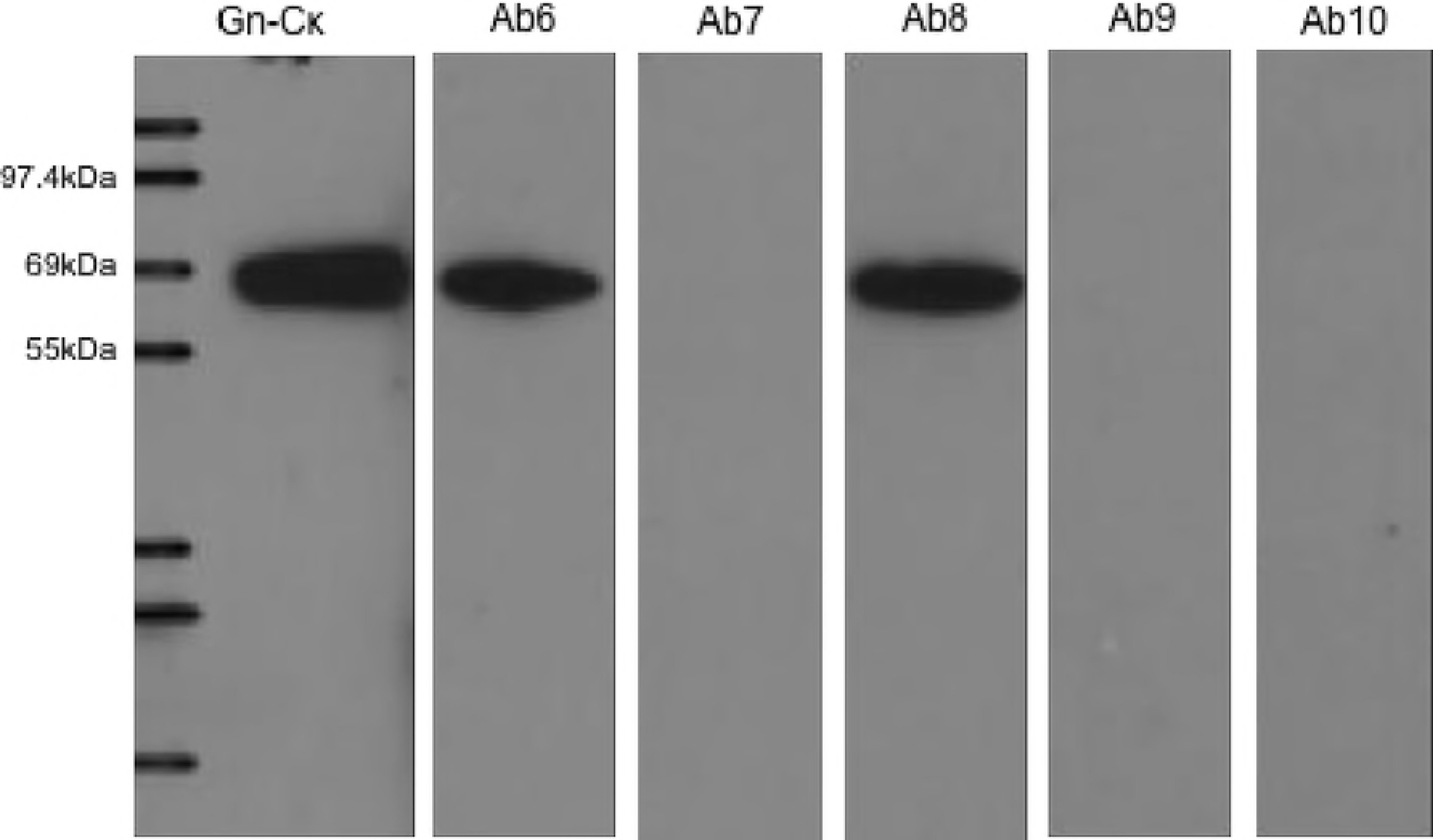
Immunoblot of recombinant Gn-Cκ fusion protein using anti-Gn antibodies. Recombinant SFTSV Gn-Cκ was prepared with sample buffer and reducing agent (NP0008 and NP0004, Invitrogen). The samples were then separated on a polyacrylamide gel (NP0321BOX, Invitrogen) by electrophoresis and transferred to a nitrocellulose membrane. After blocking with 5% (w/v) skim milk in Tris-buffered saline (pH 7.4) the membrane was incubated with 100 ng/ml of five (Ab6 to Ab10) SFTSV Gn specific antibodies in a scFv-Fc format. Gn bound antibodies were probed with HRP-conjugated anti-human IgG Fc antibody (31423, Invitrogen). To confirm the presence of Gn-Cκ protein, HRP-conjugated anti-human Ig kappa light chain antibody (AP502P, Chemicon) was used to directly detect Gn-Cκ. The blots were visualized using a chemiluminescent substrate (34578, Thermo Scientific).

**S6 Fig.**
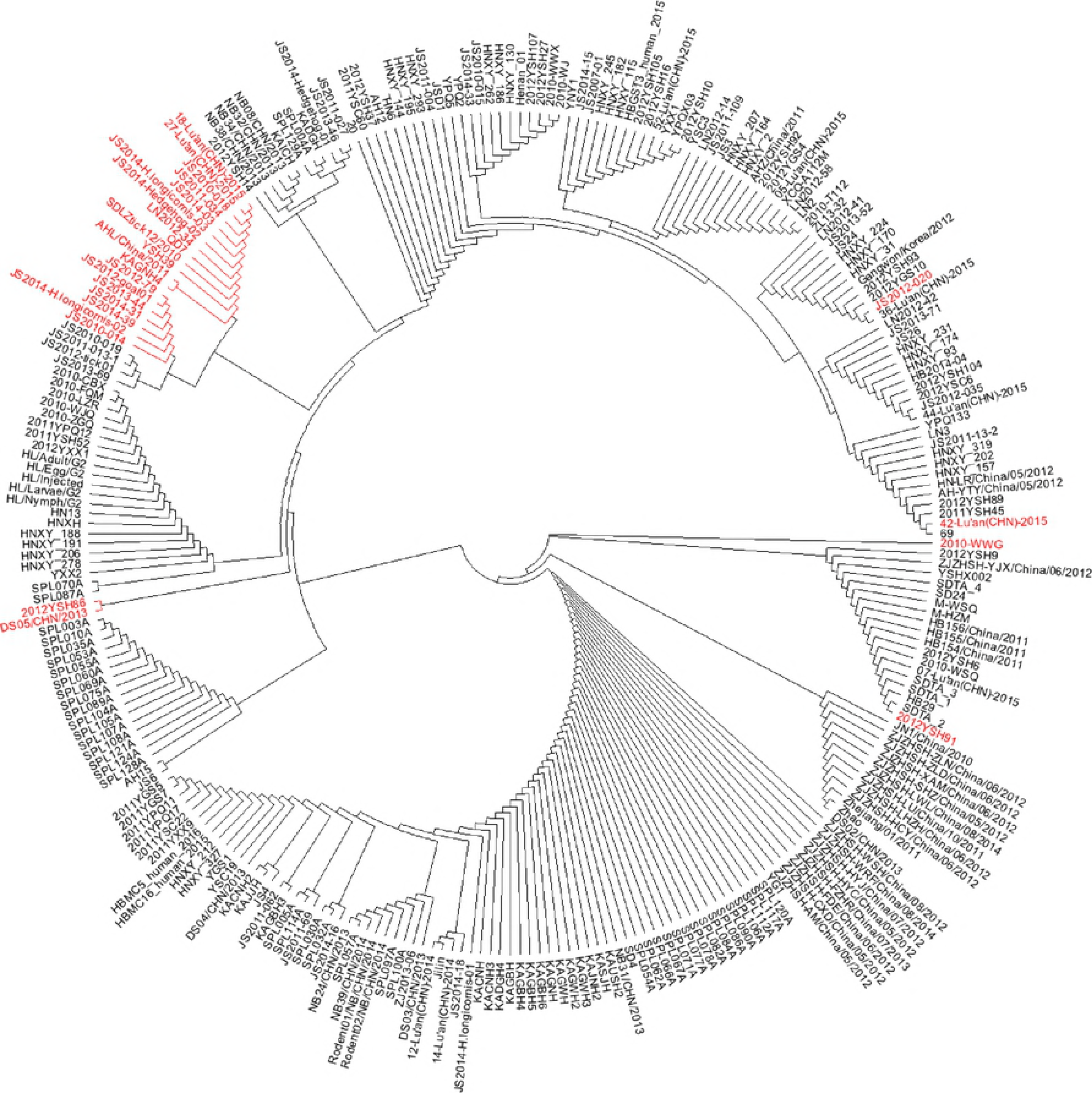
Phylogenetic analysis of sequences covering Ab10 epitope. The amino acid sequence of Gn glycoprotein from the 272 SFTSV isolates deposited in ViPR were analyzed. The sequences were trimmed to retain the amino acids from 313r389 that correspond to the residues recognized by Ab10. Trimmed sequences were analyzed and a phylogenetic tree was built in a circular tree layout using the neighbor-joining method with a Jukes-Cantor genetic distance model. The names of isolates are written beside the tip of each branch. Strain names labeled in red indicate that the Gn glycoprotein of the indicated strain is predicted to not interact with Ab10.

### Data availability

All relevant data are contained within the paper and its Supporting Information files.

